# Th2 and Th17-Associated Immunopathology Following SARS-CoV-2 Breakthrough Infection in Spike-Vaccinated ACE2-humanized Mice

**DOI:** 10.1101/2023.10.18.563016

**Authors:** Tianyi Zhang, Nicholas Magazine, Michael C. McGee, Mariano Carossino, Gianluca Veggiani, Konstantin G. Kousoulas, Avery August, Weishan Huang

## Abstract

Vaccines have demonstrated remarkable effectiveness in protecting against COVID-19; however, concerns regarding vaccine-associated enhanced respiratory diseases (VAERD) following breakthrough infections have emerged. Spike protein subunit vaccines for SARS-CoV-2 induce VAERD in hamsters, where aluminum adjuvants promote a Th2-biased immune response, leading to increased type 2 pulmonary inflammation in animals with breakthrough infections. To gain a deeper understanding of the potential risks and the underlying mechanisms of VAERD, we immunized ACE2-humanized mice with SARS-CoV-2 Spike protein adjuvanted with aluminum and CpG-ODN. Subsequently, we exposed them to increasing doses of SARS-CoV-2 to establish a breakthrough infection. The vaccine elicited robust neutralizing antibody responses, reduced viral titers, and enhanced host survival. However, following a breakthrough infection, vaccinated animals exhibited severe pulmonary immunopathology, characterized by a significant perivascular infiltration of eosinophils and CD4^+^ T cells, along with increased expression of Th2/Th17 cytokines. Intracellular flow cytometric analysis revealed a systemic Th17 inflammatory response, particularly pronounced in the lungs. Our data demonstrate that aluminum/CpG adjuvants induce strong antibody and Th1-associated immunity against COVID-19 but also prime a robust Th2/Th17 inflammatory response, which may contribute to the rapid onset of T cell-mediated pulmonary immunopathology following a breakthrough infection. These findings underscore the necessity for further research to unravel the complexities of VAERD in COVID-19 and to enhance vaccine formulations for broad protection and maximum safety.

**Significance statement:** This research investigates the safety and efficacy of a Spike protein subunit vaccine adjuvanted with Alum and CpG in an ACE2-humanized mouse model, simulating SARS-CoV-2 breakthrough infections. The study reveals that despite robust protection against severe COVID-19, vaccinated mice exhibit substantial pulmonary immunopathology, including eosinophilia and enhanced Th2 effector immunity, following breakthrough infections. Surprisingly, the study also uncovers a significant systemic Th17 inflammatory response in vaccinated mice. This research sheds light on the potential risks associated with COVID-19 vaccine breakthrough infections and the need for a comprehensive understanding of vaccine-induced immune responses, emphasizing the importance of ongoing research, surveillance, and careful vaccine development for both protection and safety in the fight against the COVID-19 pandemic.

## Introduction

Coronavirus disease 2019 (COVID-19), which is caused by severe acute respiratory syndrome coronavirus 2 (SARS-CoV-2), has stimulated global efforts of vaccine development and administration. The SARS-CoV-2 Spike protein has been used as a primary vaccine antigen, given its dominant role in mediating SARS-associated coronavirus infection (1). Spike-based vaccines are safe and effective in reducing severe COVID-19 (see Review (2)). However, protective immunity elicited by SARS-CoV-2 infection or COVID-19 vaccines may wane rapidly, leading to a rapid decrease in protective effectiveness within six months (3–5). In addition, due to the active evolution of SARS-CoV-2, new variants of concern can gain the ability to evade prior immunity, notably neutralizing antibodies, via selection against the immune epitopes (6). Under such circumstances, breakthrough infections in vaccinated individuals could occur fairly soon after vaccination (7, 8), independent of demographic factors or comorbidities (9).

Vaccine-associated enhanced respiratory diseases (VAERD) can occur in vaccinated individuals during breakthrough infections. It is well documented in humans that formalin-inactivated Respiratory Syncytial Virus (RSV) vaccines could cause lung eosinophilia, leading to enhanced respiratory diseases in infants and young children during breakthrough RSV infections (10). While VAERD has not been reported in humans following COVID-19 breakthrough infections, in animal models, SARS-associated coronavirus vaccines have been shown to enhance type 2 immunopathology (IL-4/5/13 and eosinophils) in the lung following breakthrough infections (11–13). This has been reported in multiple animal species including mice, hamsters, ferrets and non-human primates, and even more concerning, with multiple vaccine types including inactivated vaccine, virus-like particle vaccine containing multiple virus proteins, Spike-expressing viral vector vaccine, and Spike protein subunit vaccine (11–13). Although adjuvants are not required for the development of SARS-associated coronavirus VAERD in animals, aluminum-based (Alum) adjuvants can aggravate the observed vaccine-associated Th2-biased pulmonary immunopathology for Spike protein subunit vaccine (13). Alum-based adjuvants predominantly skew towards Th2 effector cell differentiation and a strong humoral immunity (14); they are also the most widely used adjuvants in human vaccines (15). CpG oligodeoxynucleotides (CpG), a soluble Toll-like receptor 9 (TLR9) ligand, can increase both cellular and humoral immunity, particularly with strong Th1 and CD8^+^ T cell immunity (16). In aged BALB/c mice, the combination of Alum and CpG has been used in COVID-19 subunit vaccines containing Spike Receptor-Binding Domain (RBD) as antigen (17, 18). Compared to Alum or CpG alone, the combination of both adjuvants enhanced host protection against a mouse-adapted SARS-CoV-2 strain (18). It is however unclear whether this combined adjuvant strategy is safe and effective in humans.

In this study, we investigated safety and efficacy of a Spike protein subunit vaccine adjuvanted with Alum and CpG in an ACE2-humanized mouse model, along with the human isolate of SARS-CoV-2 (USA-WA1/2020). As expected, vaccination with Spike protein in combination with Alum + CpG adjuvant induced robust neutralizing antibody production, Th1 effector cells, and Spike-specific CD8^+^ T cells. However, in the face of breakthrough infections, despite a significant reduction of mortality and lung viral load, vaccinated mice suffered substantial pulmonary immunopathology. As previously reported, the observed lung pathology is associated with eosinophilia and enhanced Th2 effector immunity. To our surprise, we also observed a substantial systemic Th17 response, most prominently in the lung. Our results reveal the predominant Th17-associated inflammatory response in ACE2-humanized mice, following COVID-19 vaccination and breakthrough infection. This model may be used to investigate COVID-19 vaccine-associated enhanced respiratory diseases.

## Results

### Protective effects of Spike vaccination in breakthrough infection in ACE2-humanized mice

To further investigate the protective and immunopathogenic effects of vaccine-induced host immunity in COVID-19 vaccination in a humanized animal model, we immunized K18-hACE2 mice intramuscularly with a prime-boost regimen with 5 μg per dose of SARS-CoV-2 Spike protein mixed with Alum and CpG adjuvants (Fig. 1A). Following vaccination, to simulate breakthrough infection we infected animals with increasing dosages of virus (Fig. 1A). ELISA and neutralization assays on post-vaccination but pre-exposure sera verified that our SARS-CoV-2 Spike protein vaccination induced high levels of Spike-specific IgG (see *SI Appendix* Fig. S1A) and high titers of neutralizing antibodies (Fig. 1B). Spike-protein vaccination provided protection by preventing severe morbidity and mortality (Fig. 1C). Notably, while the vaccination protected against the infection by reducing the overall viral loads in the lungs, the two-dose virus exposure allowed the establishment of a breakthrough infection in the vaccinated animals (∼10^12^ versus ∼10^8^ genome copies/gram lung tissue, comparing un-vaccinated versus vaccinated animals) (Fig. 1D).

**Fig. 1:**
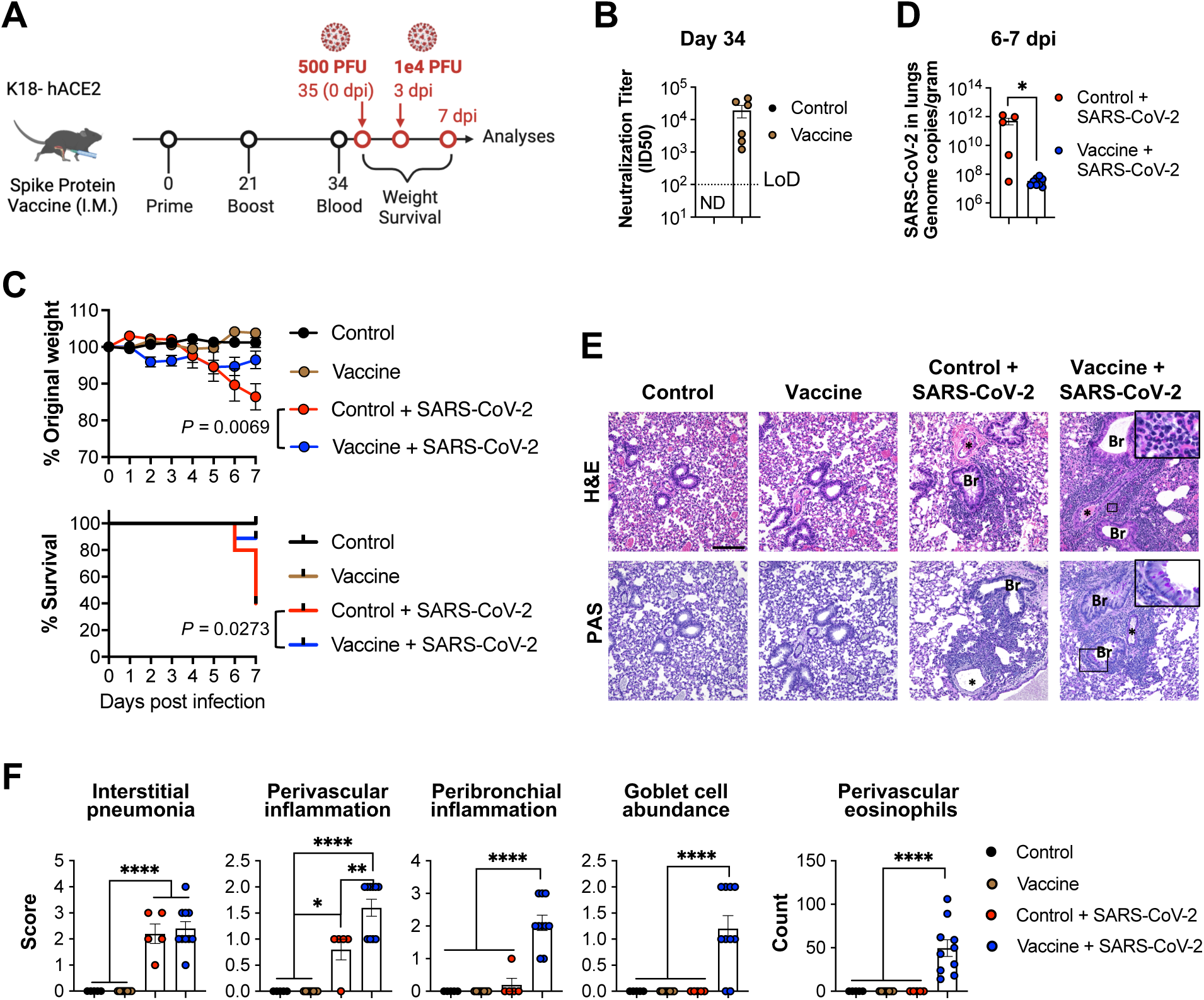
Protective effects and VAERD in hACE2 transgenic mice following SARS-CoV-2 Spike protein vaccination and breakthrough infection. (**A**) Schematic of experimental design. K18-hACE2 mice were vaccinated with PBS (Control) or SARS-CoV-2 Spike protein (Vaccine) intramuscularly (I.M.), followed by challenge with increasing dosage of SARS-CoV-2 administered intranasally (I.N.). (**B**) Neutralizing antibody titers following vaccination prior to infection. Prior to infection (day 34 post vaccine priming), sera were collected from mice and analyzed in neutralization assays to verify the induction of neutralizing antibodies by vaccination (Control n = 5; Vaccine n = 6 randomly sampled from each cage). Limit of detection, LoD = 100. (**C**) Percentage of original body weight and survival at the indicated time points. *P* value in weight curve was calculated by two-way ANOVA. *P* value in survival curve was calculated by log-rank test. Animals that lost ≥ 20% body weight were euthanized and recorded as dead, and organs were collected for analysis. All animals were euthanized at the beginning of day 7 post infection. (**D**) SARS-CoV-2 viral load (genome copies/gram) in lungs (Control + SARS-CoV-2 n = 5; Vaccine + SARS-CoV-2 n = 10). * *p* ≤ 0.05 by two-tailed student’s *t* test. (**E**) Representative images of H&E and PAS staining of lung tissue sections. Magnification: 200 ×. Scale bar = 100 μm. Br denotes bronchioles and * denotes pulmonary vasculature. Rectangles indicate zoom-in areas depicting increase in perivascular eosinophils and higher frequency of goblet cells in the bronchiolar epithelium. (**F**) Histopathological scores. Control n = 3; Vaccine n = 8; Control + SARS-CoV-2 n = 5; Vaccine + SARS-CoV-2 n = 10. \**p* < 0.05, ** *p* < 0.01, **** *p* < 0.0001 by one-way ANOVA with multiple comparisons. Data presented as Mean ± S.E.M.

### Type 2 pulmonary immunopathology following Spike vaccination and breakthrough infection

Despite a significantly improved survival rate, vaccinated animals appeared distressed assuming a hunched posture, exhibiting reluctance to move and lethargy. Histological staining of the lung sections revealed that the breakthrough infections caused a significant level of interstitial pneumonia comparable to that observed in the un-vaccinated and infected mice (Fig. 1E & F). Moreover, breakthrough infections in vaccinated animals resulted in elevated levels of perivascular and peribronchial inflammation, as compared to the lungs of the un-vaccinated and infected mice (Fig. 1E & F). The observed severe immunopathology in vaccinated mice following breakthrough infections was associated with significant increase in 132 the abundance of goblet cells and increased counts of perivascular eosinophils (Fig. 1E & F). Flow cytometric analyses of cells recovered from the lungs further revealed significant accumulation of eosinophils and IgE-bound basophils/mast cells in the vaccinated mice with breakthrough infections (Fig. 2A & B). We also noticed that, prior to infection, vaccination with Spike protein in Alum and CpG adjuvants was sufficient in inducing high levels of IgE production detected in the sera collected 7 days post the vaccine boost (see *SI Appendix* Fig. S1B). Also, we detected a trend of higher levels of Spike specific IgE production in the vaccinated mice (see *SI Appendix* Fig. S1B). On the other hand, the increased numbers of interstitial macrophages and dendritic cells in the lungs following SARS-CoV-2 infection were not affected by vaccination, while the infection-driven neutrophil enrichment in the lung was significantly reduced by vaccination, though not to levels comparable to those observed in un-infected mice (Fig. 2B). In agreement with previous findings (13), our results show that immunization with Spike protein followed by a breakthrough SARS-CoV-2 infection causes a type 2-associate inflammatory response, prominently eosinophilia in the lung, which leads to vaccine-associated respiratory pathology in animal models of COVID-19.

**Fig. 2:**
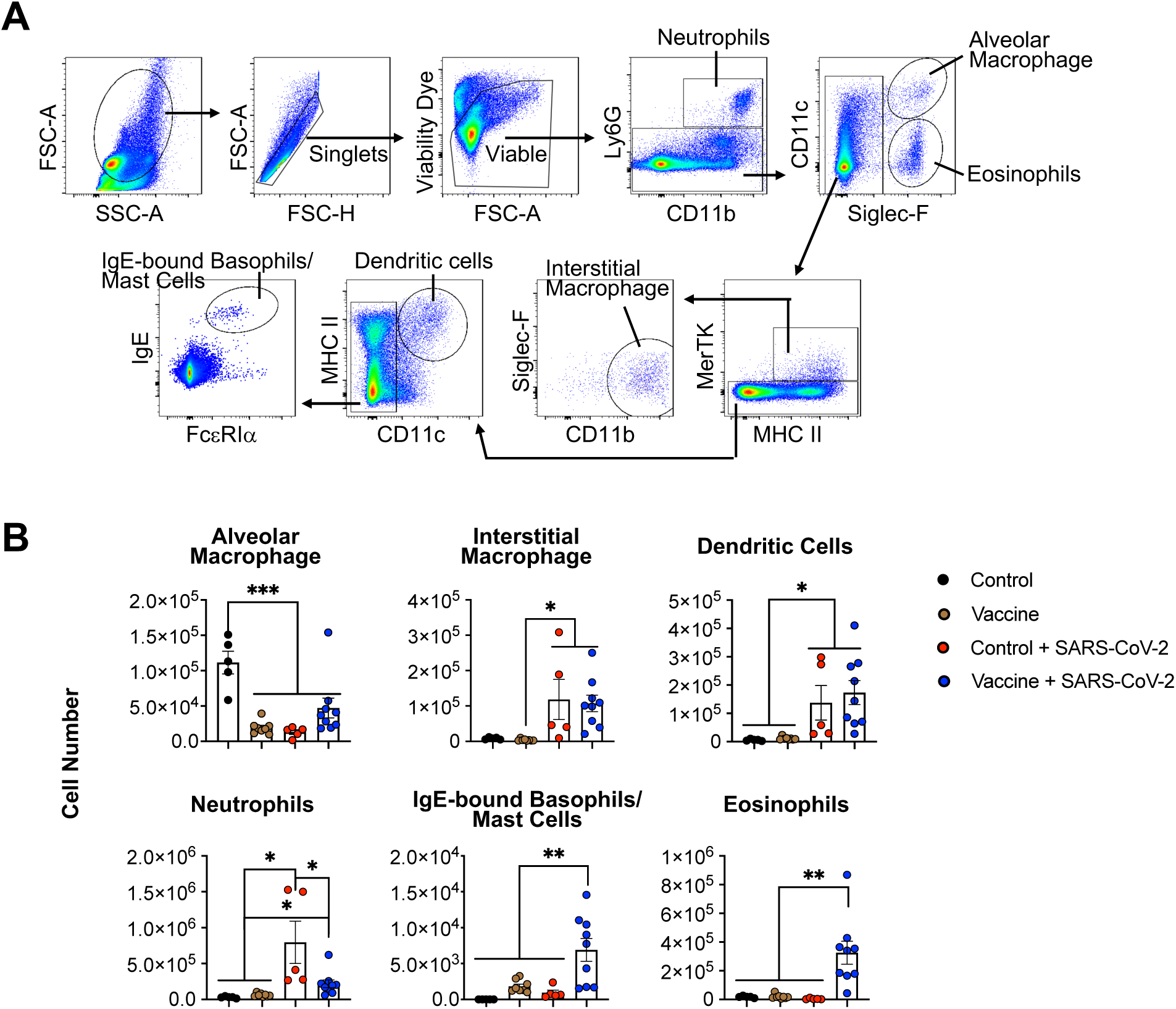
Pulmonary innate immune cell profile. (**A**) Representative flow cytometric plots for innate immune cell gating. Cells were stained and fixed overnight before analyses. (**B**) Numbers of the indicated innate immune cells in the lungs. Control n = 5; Vaccine n = 8; Control + SARS-CoV-2 n = 5; Vaccine + SARS-CoV-2 n = 9. \**p* < 0.05, ** *p* < 0.01, *** *p* < 0.001 by one-way ANOVA with multiple comparisons. Data presented as Mean ± S.E.M.

### Massive lung infiltration of CD4^+^ T cells following Spike protein vaccination and breakthrough infection

We investigated the effects of Spike vaccination on lymphocytes in the lungs of the mice with breakthrough infections by flow cytometric analyses (Fig. 3A). In the absence of an infection, immunization with Spike protein did not cause lymphocyte enrichment in the lungs (Fig. 3B). However, following infection, the numbers of NK cells, *invariant* NKT cells, and CD8+ T cells in the lungs substantially increased, with no significant differences between un-vaccinated and vaccinated animals (Fig. 3B). In contrast, we observed significantly 155 enhanced lung infiltration of B cells, γδ T cells and CD4^+^ T cells in the vaccinated and infected animals (Fig. 3B), with the lung-infiltrating CD4^+^ T cells representing the most abundant lymphocyte population in the vaccinated animals with a breakthrough infection (Fig. 3B). To further determine the histological locations of CD4^+^ T cell infiltration, we subjected lung tissue sections to immunohistochemical analysis, and observed CD4^+^ T cells throughout the lung tissue sections with a perivascular enrichment (Fig. 3C). These data suggest that these CD4^+^ T cells might be recruited to the lungs during breakthrough infection

**Fig. 3:**
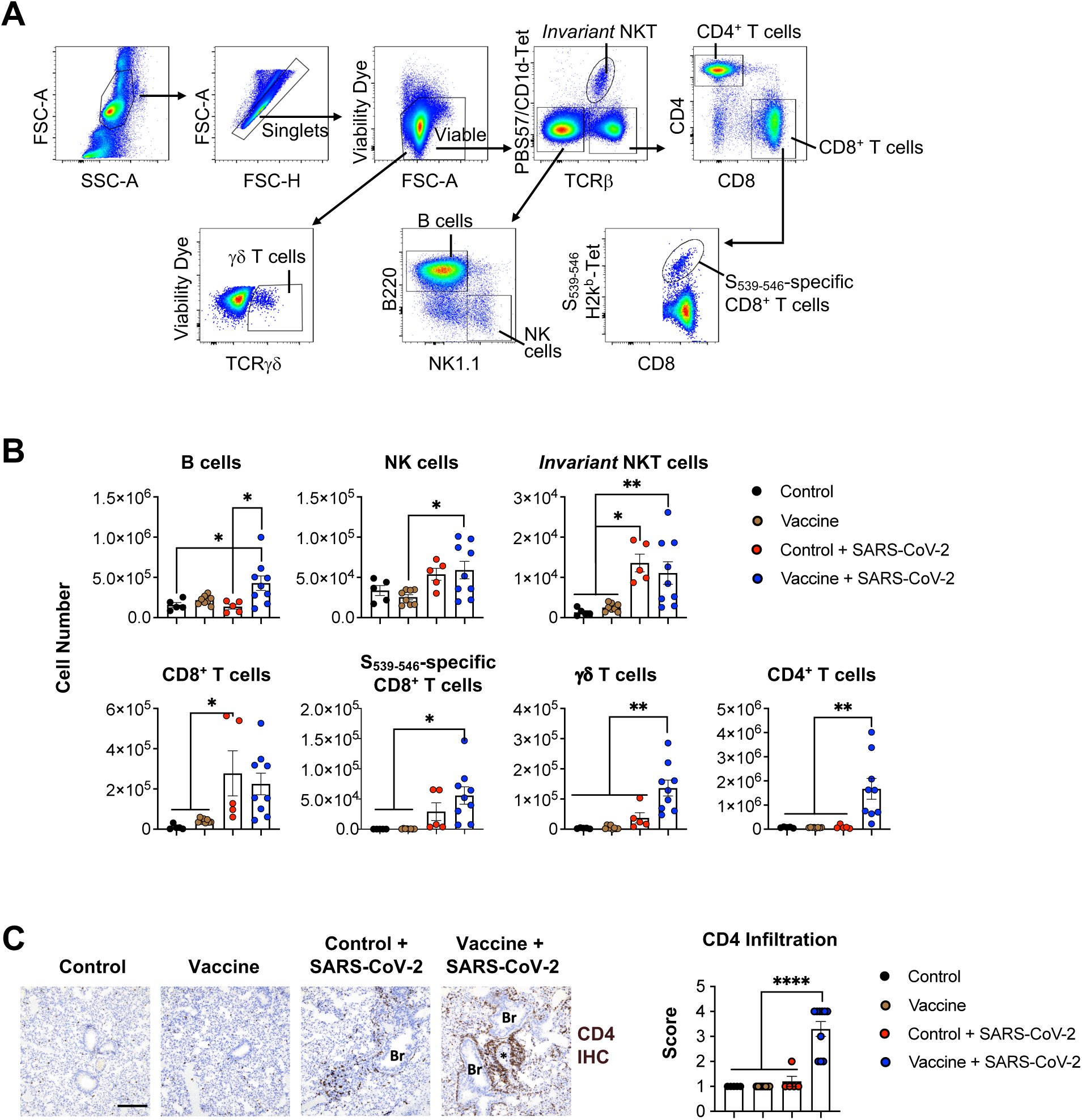
Pulmonary lymphocyte profile. (**A**) Representative flow cytometric plots for lymphocyte gating. Cells were stained and fixed overnight before analyses. (**B**) Numbers of the indicated lymphocytes in the lungs. (**C**) Representative immunohistochemical staining of CD4 (brown) and scores of CD4^+^ T cell infiltration in lung tissue sections. Magnification: 200 ×. Scale bar = 100 μm. Br denotes bronchioles and * denotes pulmonary vasculature. Control n = 5; Vaccine n = 8; Control + SARS-CoV-2 n = 5; Vaccine + SARS-CoV-2 n = 9. \**p* < 0.05, ** *p* < 0.01, *** *p* < 0.001 by one-way ANOVA with multiple comparisons. Data presented as Mean ± S.E.M.

### Th2 and Th17-associated pulmonary inflammatory cytokine responses following Spike vaccination and breakthrough infection

Given the type 2-associated pulmonary immunopathology and the massive CD4^+^ T cell infiltration in the lungs of ACE2-humanized animals following Spike immunization and breakthrough infection (Fig. 1-3), we suspected that there was an enhanced Th2 cytokine response involved. To further compare the levels of other T helper subtype-associated cytokine production, we quantified the levels of gene expression of cytokines associated with different T helper cell subtypes (19, 20), including *Ifng* (Th1), *Il4/Il13* (Th2), *Il17a* (Th17), *Il21* (Tfh), *Csf2* (encodes GM-CSF), and *Il10* (Regulatory T cells) (Fig. 4). We found that SARS-CoV-2 infection induced high levels of expression of *Ifng* in both un-vaccinated and vaccinated mice, and as expected, vaccination with Spike protein followed by a breakthrough infection led to elevated expression of Th2 cytokines *Il4* and *Il13*. Similarly, the expression of *Il21* was significantly higher in the vaccinated and infected animals compared to untreated controls or mice that were vaccinated but not infected. There was a trend of increased expression of *Il21* in the vaccinated and infected group as compared to the un-vaccinated and infected mice, suggesting a strong Tfh cell response in the vaccinated and infected animals. Vaccination showed an inhibitory effect on virus-associated induction of the expression of *Il10* and *Csf2.* To our surprise, the expression of *Il17a* was significantly higher in the vaccinated and infected animals, as compared to any other groups, suggesting that a Th17-associated inflammatory response was likely involved in the development of the VAERD observed in the Spike-vaccinated mice that suffer a breakthrough infection (Fig. 4).

**Fig. 4:**
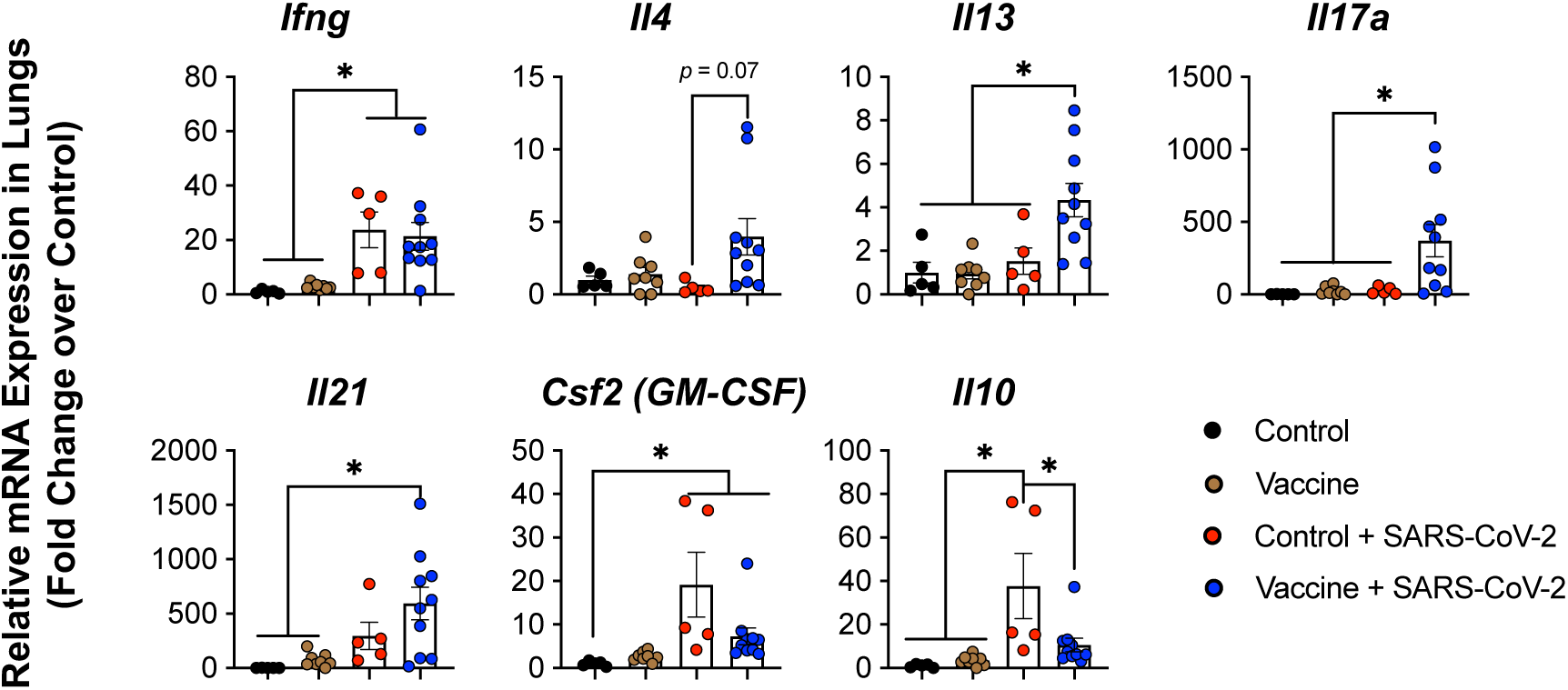
Pulmonary cytokine transcript production. Relative fold change expression of mRNAs encoding IL-4, IL-5, or IL-13 was determined using quantitative RT-PCR. After normalization to *Gapdh* expression, fold changes were calculated by normalization to the averages in the Control group. Control n = 5; Vaccine n = 8; Control + SARS-CoV-2 n = 5; Vaccine + SARS-CoV-2 n = 10. \**p* < 0.05 by one-way ANOVA with multiple comparisons. Data presented as Mean ± S.E.M.

### A systemic Th17 inflammatory response following Spike vaccination and breakthrough infection

We suspected that the CD4^+^ T cell population was responsible for the elevated *Il17a* expression, given that these were the most abundant infiltrating lymphocytes in the lungs of animals with severe VAERD. To confirm this hypothesis, we stimulated cells recovered from the lung, draining lymph node of the vaccinated site, and the spleen with a cell stimulation cocktail (containing PMA, Ionomycin, and Brefeldin A), and performed intracellular cytokine staining for Th1 effector associated cytokines (IFN-γ and TNF-α), Th2-associated cytokine (IL-4), and Th17 cytokine (IL-17A). We found that the percentage of cells expressing Th1, Th2 or Th17 cytokines were all significantly higher in the lungs and draining lymph nodes of the vaccinated and infected group compared to mice under any of the other conditions (Fig. 5A & B). Strikingly, the increase of Th17 cell numbers in the vaccinated and infected animals appeared to be systemic, as it was highly significant in cells recovered from the spleen of the mice with VAERD (Fig. 5A & B, Spleen panels). Note that the number of γδ T cells was also significantly elevated in the vaccinated and infected mice compared to all other groups (Fig. 3B). Considering that γδ T cells are capable producers of IL-17 which can initiate a Th17 inflammatory response (21), we analyzed the production of different Th cell-associated cytokines in lung γδ T cells as well. However, we did not find evidence of VAERD-associated patterns of cytokine production by the γδ T cells (see *SI Appendix* Fig. S2). Our results suggested that the COVID-19 VAERD observed in the ACE2-humanized mouse model is associated with enhanced local Th1, Th2 and Th17 inflammatory responses, as well as systemic Th17-associated inflammation.

**Fig. 5:**
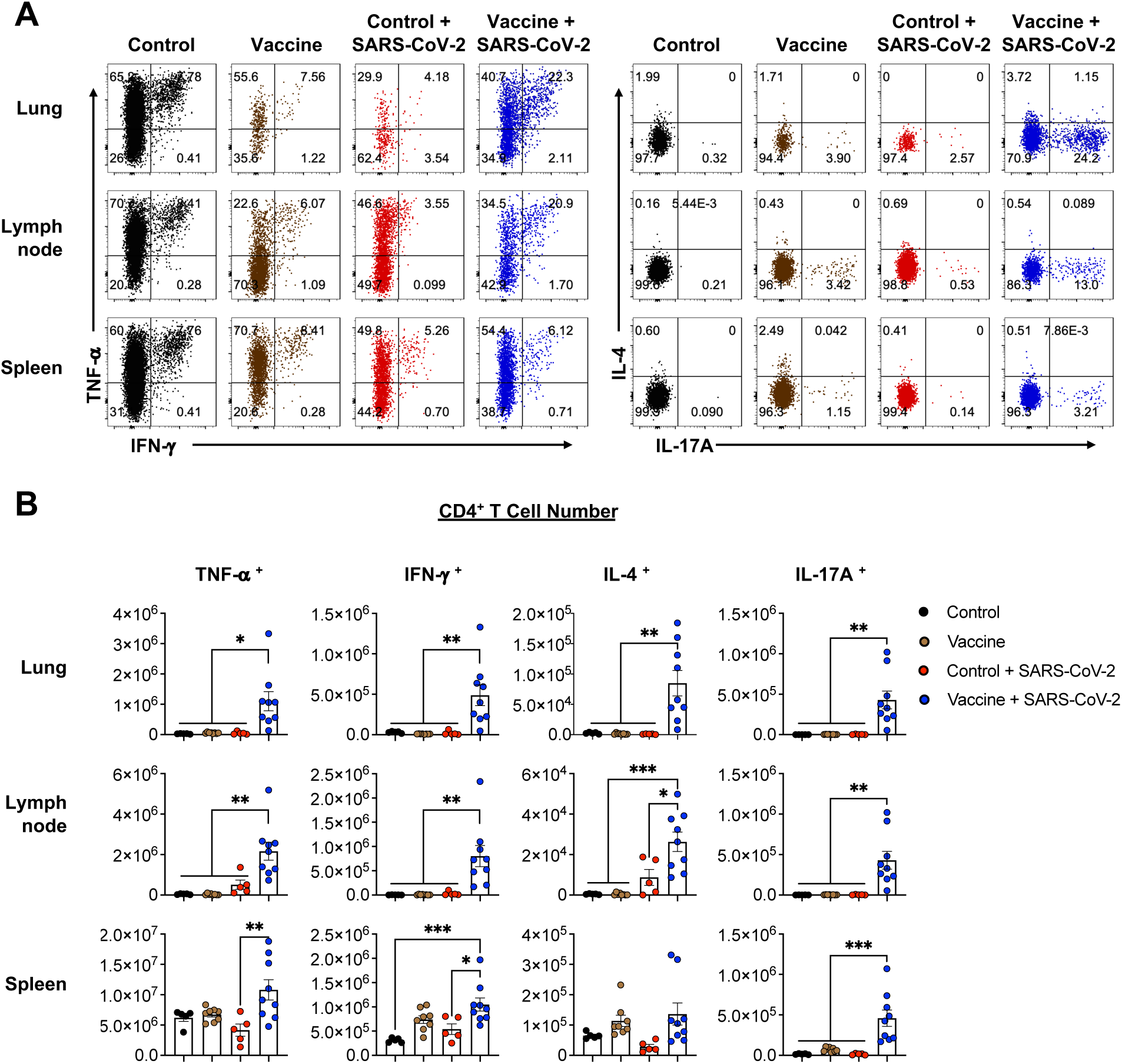
Effector cytokine production by CD4^+^ T cells. (**A**) Representative flow cytometric plots of TNF-α, IFN-γ, IL-4 and IL-17A expression by viable CD4^+^ T cells in lung, lymph node and spleen. (**B**) Numbers of CD4^+^ T cells in lung, lymph node and spleen expressing TNF-α, IFN-γ, IL-4 and IL-17A. Control n = 5; Vaccine n = 8; Control + SARS-CoV-2 n = 5; Vaccine + SARS-CoV-2 n = 9. \**p* < 0.05 by one-way ANOVA with multiple comparisons. Data presented as Mean ± S.E.M.

## Discussion

It is undeniable that SARS-CoV-2 Spike-based vaccination is safe and effective in preventing severe and fatal COVID-19. Immunization against Spike protein using mRNA, viral vector-based or protein subunit vaccines can elicit protective immunity and prevent severe COVID-19 illness (22). Our results further support the global consensus on the protective effects of vaccines. However, while COVID-19 vaccines have been successful in reducing the severity of the disease, additional efforts are needed to enhance our understanding of the complex dynamics of COVID-19 vaccination and the potential risks associated with breakthrough infections, especially concerning the emergence of VAERD. Multiple studies in different animals models of SARS-associated coronaviral infections reported that VAERD is associated with a Th2-biased inflammatory response (11–13). Here we systematically investigated the potential risks and underlying mechanisms of immunopathogenesis of VAERD in COVID-19 breakthrough infections. Using an ACE2-humanized mouse model of COVID-19, we observed a similar Th2-associated lung immunopathology following Spike immunization and SARS-CoV-2 breakthrough infection. Furthermore, our results suggest a prominent role of Th17 inflammatory response in correlation with severe COVID-19 VAERD. To the best of our knowledge, this is the first report of a prominent Th17 inflammatory response in VAERD in animal models of SARS-associated coronaviral infections.

An increased production of Th1-associated cytokines (e.g. IFN-γ) is correlated with reduction in COVID-19 severity (23), and vaccine-induced Th1 cell responses correlate with the protective effects of the current mRNA vaccines (24). CpG ODNs that can stimulate TLR9 and trigger strong Th1 type immunity have been considered as adjuvants for vaccinations against viral and bacterial infections, as well as cancer (16). Classic Alum adjuvants are the first licensed and most widely used vaccine adjuvants. Alum adjuvants can prime a Th2-biased T cell differentiation and promote humoral responses which provide effective protective immunity against many pathogens (14). However, in severe coronavirus diseases in humans, Th2 cytokine polarization has been associated with disease severity and increased mortality (25). Additionally, Th2-skewing vaccination has consistently been linked to type 2 inflammatory immunity, notably eosinophilia, in cases of VAERD in animal models of coronaviral diseases (11–13, 26, 27). In mouse models of allergic airway inflammation, where ovalbumin (OVA) is used to sensitize mice towards airway hypersensitivity upon respiratory exposure to OVA, Alum has been shown to induce a Th2-biased immunity accompanied by the induction of IgE and eosinophils. Interestingly, the effects of Alum in the OVA model could be reverted to basal levels by co-administration of CpG (28). Similarly, the inclusion of Th1-skewing adjuvants in SARS-CoV-1 inactivated or Spike protein subunit vaccines attenuated the eosinophilic respiratory inflammatory responses during infections (29, 30). Surprisingly, in SARS-CoV-2 Spike immunization followed by breakthrough infections, we observed elevated levels of lung immunopathology, eosinophilia, IgE, and Th2 responses, even if CpG and Alum were both used in vaccination. Moreover, we observed a systemic Th17 proinflammatory response. This is likely due to the intrinsic antigenic properties of the SARS-CoV-2 Spike protein. For example, the lethal Th2-associated immunopathology following SARS-CoV-1 Spike immunization and viral infection can be induced by certain regions of the SARS-CoV-1 Spike protein such as S_597-603_ (LYQDVNC, an epitope downstream of the C terminus of the RBD) (12), which is identical to SARS-CoV-2 S_611-617_ (31). Since antibodies to this peptide have been shown to cause antibody-dependent enhancement (ADE) of infection in SARS-CoV-1, it is predicted that SARS-CoV-2 S_611-617_ may be involved in vaccine-associated enhanced pathology during infections through type 2 inflammation and potentially ADE (31). It is possible that other regions in SARS-CoV-2 Spike protein are associated with a Th17 cell priming under the Alum/CpG combined adjuvant condition.

The induction of a systemic Th17 inflammatory response during breakthrough SARS CoV-2 infection is particularly concerning. IL-17 exhibits multifaceted functions during viral infections. For example, during influenza infection, on one hand IL-17 promotes antiviral immunity and protects the hosts from severe viral disease, while on the other hand it contributes to the pathogenesis of immunopathology, lung injury, and post influenza superinfection (32). While the potential protective role of IL-17 in COVID-19 is unclear, there are documented associations between high IL-17 levels and COVID-19 pathogenesis. In severe COVID-19 patients, there are elevated levels of serum IL-17A and circulating Th17 cells (33, 34). High serum levels of IL-17 were also observed in long-COVID (35). IL-17 can not only cause skin and lung tissue inflammation and fibrosis (36, 37), but also induces mucus production and goblet cell metaplasia in lung epithelial cells (38, 39). IL-17 and neutrophils are also linked to thrombus formation in acute myocardial infarction (40). Combined with TNF-a, IL-17A moderates thrombus growth on endothelial cells *in vitro* (41). In mouse models, IL-17A promotes deep vein thrombosis by enhancing platelet activation/aggregation, neutrophil infiltration, and endothelial cell activation (42). These features of Th17-associated inflammatory diseases should be considered in the analysis of potential COVID vaccine-associated adverse effects.

In conclusion, this research underscores the complexity of COVID-19 vaccination and the need for a comprehensive understanding of vaccine-induced immune responses. While vaccines remain a vital tool in combating pandemics, the potential for VAERD highlights the importance of ongoing research, surveillance, and careful vaccine development to achieve broad protection and maximal safety.

## Materials and Methods

### Mice

B6. Cg-Tg(K18-ACE2)2Prlmn/J (K18-hACE2; Strain #:034860) mice were purchased from the Jackson Laboratory and housed in the Division of Laboratory Animal Medicine, at Louisiana State University School of Veterinary Medicine. All experiments were approved by the Institutional Animal Care and Use Committee at Louisiana State University.

### Immunization and SARS-CoV-2 infection

Male mice at the age of around 10 weeks were immunized intramuscularly (I.M., ipsilateral administration) in a prime-boost regimen (Fig. 1, day 0 and day 21) with 5 μg/dose of recombinant SARS-CoV-2 Spike protein (S1+S2 extracellular domain) (Bon Opus Biosciences, #BP040) adjuvanted with 100 μg/dose alhydrogel adjuvant 2% (InvivoGen, #vac-alu-250) and 20 μg/dose CpG ODN 2395 – TLR9 ligand (InvivoGen, #tlrl-2395-5) in 100 μL of PBS (pH 7.4). Mice vaccinated with PBS were used as negative controls. Blood was collected from the facial vein at the indicated time points, prior to infection, to confirm the induction of Spike-specific antibodies, neutralizing antibodies, and total IgE. Control and vaccinated mice were initially challenged with a low dose (500 PFU) of SARS-CoV-2 (USA-WA1/2020, BEI Resources, # NR-52281), followed by a high dose (10,000 PFU) infection of the same virus 3 days later. Viruses were delivered to mice intranasally in 50 μL of sterile PBS. Infected animals were monitored daily. Mice that did not respond to back padding or lost ≥ 20% of original weight were humanely euthanized and recorded as a death event. Seven days post the initial infections (dpi), all viable animals were euthanized for analyses. Serum, spleen, lung and mediastinal lymph nodes were collected.

### Serum neutralizing antibody titration

Calu-3 cells (ATCC; HTB-55) (1×10^5^/well in 50 μL) were seeded in 96-well plates, followed by incubation in 37 °C, with 5% CO_2_, and 6-12 h later, cells were infected with SARS-CoV-2 (USA-WA1/2020; 1000 PFU). Titrated sera were mixed with virus in 50 μL and incubated in 37 °C for 30 minutes, and then applied to the seeded cells. Forty hours post infection, supernatant was removed, and cells were incubated with 60 μL of 0.25% trypsin-EDTA (25200072; ThermoFisher Scientific) for 2-3 mins, followed by the addition of 120 μL of 6% paraformaldehyde (PFA)/PBS. Cells were then transferred to V-bottom plates and fixed overnight. Fixed cells were then permeabilized and stained (using BioLegend Buffer 421002) for intracellular SARS-CoV-2 nucleocapsid (N) protein using a monoclonal Alexa Fluor 647-conjugated anti-N antibody (kind gift from Dr. Yongjun Guan). Flow cytometric data was acquired with an automatic plate reader system (CytoFlex, Beckman Coulter, Inc.), and analyzed using FlowJo Software (version 10.1, FlowJo, LLC). Percentages of infection and dilution factors were used for nonlinear regression to calculate for doses for 50% inhibition (ID50) indicated by the dilution factors in Prism 9.3.1 (GraphPad).

### Quantitative reverse-transcription PCR (qRT-PCR)

The post-caval lobe of the lung was weighed and homogenized in 1 mL of PBS (pH 7.4) using a TissueLyser LT (Qiagen). RNA was extracted using TRIzol™ LS Reagent (Invitrogen) and cDNA was generated using Protoscript II First Strand cDNA Synthesis Kit (New England Biolabs). SARS-CoV-2 was detected by PCR using the 2019-nCoV RUO Kit (IDTDNA, # 10006713). To quantify viral genome copies, quantitative PCR (qPCR) Extraction Control from Heat-Inactivated SARS-Related Coronavirus 2, Isolate USA-WA1/2020 (BEI Resources, # NR-52347) was used to generate the standard curve, which was used for calculating the genome copies per gram in the lung samples. To quantify cytokine gene transcript levels, “Best Coverage” gene probes for mouse Gapdh (Mm99999915_g1), Ifng (Mm01168134_m1), Il4 (Mm00445259_m1), Il13 (Mm00434204_m1), Il17a (Mm00439618_m1), Il21 (Mm00517640_m1), Gmcsf (Mm01290062_m1), and Il10 (Mm01288386_m1) (Applied Biosystems TaqMan Assays) were used as per manufacturer’s instructions. Relative mRNA expression levels were calculated by normalizing cytokine gene values to the internal loading control Gapdh, and normalized against the average of cytokine levels in the control (un-immunized, un-infected).

### Histopathology and immunohistochemistry

Sections of the lung were fixed in 10% neutral buffered formalin for a minimum of 72 hours and subsequently processed and embedded in paraffin following standard procedures. Serial four-micron tissue sections were either stained with hematoxylin and eosin (H&E) or subjected to the Periodic acid Schiff reaction (PAS) following standard procedures.

For mouse CD4 immunohistochemistry, sections were mounted on positively charged Superfrost Plus slides (Fisher Scientific) and processed using the automated BOND-RXm system (Leica Biosystems). Following automated deparaffinization, tissue sections were subjected to automated heat-induced epitope retrieval using a ready-to-use EDTA-based solution (pH 9.0; Leica Biosystems) for 20 min at 100°C. Subsequently, endogenous peroxidase was quenched by incubation with 3% hydrogen peroxide for 5 mins, followed by incubation with a recombinant rabbit anti-mouse CD4 monoclonal antibody (clone EPR19514; Abcam) diluted 1:4,000 in a ready-to-use antibody diluent (Leica Biosystems) for 30 mins at room temperature. Sections were then washed and stained with a polymer-labeled goat anti-rabbit IgG conjugated to HRP for 8 mins, followed by incubation with HRP substrate for 10 mins, at room temperature. Counterstaining was performed with hematoxylin for 5 min and slides were finally mounted with Micromount (Leica Biosystems).

Scoring of stained tissue sections was based on a cumulative score (maximum attainable score of 17) derived from 5 specific histologic features evaluated in 5 random 20 × fields following the scoring rubrics shown in *SI Appendix* Table S1. Additionally, absolute eosinophil counts within perivascular cuffs were performed in 3 randomly selected perivascular cuffs.

### Flow cytometric analyses

For flow cytometric analyses, spleens, mediastinal lymph nodes, and lungs were homogenized via grinding against strainers, followed by filtering through the strainers. Red blood cells were lysed with RBC Lysis Buffer (Tonbo Biosciences, #TNB-4300-L100), followed by washing and pelleting. Cells were resuspended in full RPMI-1640 media for counting, staining and analysis via flow cytometry. Surface protein staining was performed with antibodies for surface markers in PBS pH 7.4, in the presence of Fc Block (TruStain FcX™, BioLegend, # 101319) and fixable viability dye (Ghost Dye™ Violet 510, Tonbo Biosciences) at room temperature for 30 mins. To measure cytokine production after *ex vivo* stimulation, 500 μL single-cell suspensions were incubated at 37 °C for 6 h with 5% CO_2_ in 48-well plates in the presence of Cell Stimulation Cocktail (Tonbo Biosciences, #TNB-4975). Stimulated cells were stained with surface markers as described above, then were fixed with 4% PFA (diluted by PBS) at 4 °C overnight, followed by permeabilization and staining using the eBioscience Intracellular Permeabilization Buffer (Invitrogen, #88-8824-00). Flow data was acquired using the LSRFortessa X-20 cell analyzer (BD Bioscisence) and analyzed using FlowJo. The list of antibodies used is available in *SI Appendix*.

### Statistical analysis

T-test, one-way ANOVA and two-way ANOVA results with *p* ≤ 0.05 were considered statistically significant. Survival rates were compared by log-rank test, with *p* ≤ 0.05 considered as significantly different. All the statistical tests were performed using GraphPad Prism version 9.3.1 (GraphPad, San Diego, CA).

## Acknowledgements and funding sources

The SARS-CoV-2 USA-WA1/2020 strain and its quantitative PCR standard control were from the BEI Resources, while the CD1d and H2kb mouse tetramers were from the NIH Tetramer Core Facility. We also thank Dr. Yongjun Guan (previously at Antibody Biopharm Inc.) for his kind gift of the Alexa Fluor 647-labeled anti-SARS-CoV-2 N antibody, members of the Huang and Veggiani groups for helpful discussions. This research was supported in part by grants from the National Institutes of Health (P20 GM130555-6610, P20 GM130555-5011 and R01 AI151139) and a Big Idea Research Grant from the Provost’s Fund at Louisiana State University.

## Data availability statement

Datasets are available upon reasonable request.

## Authors’ contributions

T.Z., N.M., M.C.M., M.C. and W.H. performed experiments and analyzed data; T.Z. and W.H. designed experiments and wrote the manuscript; W.H. secured funding and supervised the research; all authors reviewed, edited and finalized the manuscript.

