## Supplemental Materials for "Th2 and Th17-Associated Immunopathology Following SARS-CoV-2 Breakthrough Infection in Spike-Vaccinated ACE2-humanized Mice"

### Methods

***Antibody detection by enzyme-linked immunosorbent assay (ELISA).***

The concentrations of serum total IgE were measured by ELISA using the Mouse IgE ELISA Set (BD Biosciences, #555248) as per manufacturer instructions. To measure the Spike-specific IgG and IgE, Spike protein (BioTime, #P5008H1-100) was coated on High Binding UltraCruz ELISA Plates (Santa Cruz Biotechnology, #sc-204463) for 24 to 96 h at 4°C (1 -2 μg/mL in 50 -100 μL PBS, pH 7.2). Serum samples were tested for Spike-specific IgG in 30,000-fold dilution as previously reported (1, 2). To detect anti-Spike IgE, coated plates were washed with PBST (0.1% TWEEN 20 in PBS) and blocked with 2% non-fat milk in PBST for 2 h at room temperature (RT). Serum samples with 25-fold dilution in assay buffer (1% non-fat milk in PBST) were used for primary incubation (100 μL/well) for 1.5 ~ 2 hours at RT. After stringent plate washing, 100 μL of the HRP-conjugated Rat anti-mouse IgE detection antibody (3) (clone 23G3; Southern Biotechnology Associates, Inc., #1130-05) were added to each well (1:1000 in assay buffer) and incubated at RT for 1 hour. Following washing, 3, 3′,5,5′-Tetramethylbenzidine (TMB) liquid Substrate (Sigma-Aldrich, #T0440) and TMB Stop solution (2N sulfuric acid) were used for detection of HRP activity. Plates were read in a microplate ELISA reader at 450 and 620 nm.

***Fluorescent antibodies used (listed in the format of "Fluorophore-target (clone)")***

Pacific Blue (PB)-Ly6G (1A8), FITC-IgE (RME-1), PE-SiglecF (S17007L), Alexa Fluor 700-CD11b (M1/70), PerCP/Cy5.5-I-A/I-E (MHC II) (M5/114.15.2), PE-Cy7-FcεRIα (MAR-1), APC-CD4 (GK1.5), Alexa Fluor 700-B220 (RA3-6B2), PE-Cyanine7-NK1.1 (PK136), APC-Cy7-TCRβ (H57-597), FITC-IL-2 (JES6-5H4), PE/Dazzle 594-IL-17A (TC11-18H10.1), APC-IL-4 (11B11), and PE/Cy7-IFN-γ (XMG1.2) were from BioLegend. APC-Cy7-CD4 (GK1.5),

PerCP/Cy5.5-CD8α (53-6.7), PE-Cy7-CD8α (53-6.7), PE-Cy7-NK1.1 (PK136) were purchased from Tonbo Biosciences. Super Bright 600-MerTK (DS5MMER), APC-eFluor 780-CD11c (N418), eFluor 450-TCRγ/δ (GL-3), and Alexa Fluor 700-TNFα (MAb11) were from ThermoFisher Scientific. The following reagents were obtained through the NIH Tetramer Core Facility: mouse CD1d tetramer loaded with PBS-57 (Brilliant Violet 421, #57065), H-2K(b) tetramer loaded with SARS-CoV-2 $S_{539-546}$ (VNFNFNGL; PE, #57074).

**Supplemental Table S1. Scoring Rubrics of Lung Histopathology.**

| Parameter evaluated | Score | Interpretation |
|---|---|---|
| **Interstitial pneumonia** | 0 | None |
| | 1 | <10% of parenchyma affected; minimal |
| | 2 | 10-25% of parenchyma affected; mild |
| | 3 | 25-50% of parenchyma affected; moderate |
| | 4 | >50% of parenchyma affected; severe |
| **Peribronchiolar inflammation** | 0 | None |
| | 1 | Single layer of inflammatory cells |
| | 2 | 2-3 layers of inflammatory cells |
| | 3 | >3 layers of inflammatory cells |
| **Perivascular inflammation** | 0 | None |
| | 1 | Cuffs of approximately <50 microns thick |
| | 2 | Cuffs of approximately 50-100 microns thick |
| | 3 | Cuffs of approximately >100 microns thick |
| **Goblet cell hyperplasia** | 0 | No visible goblet cells |
| | 1 | Few noticeable goblet cells |
| | 2 | Frequent goblet cells |
| | 3 | Diffuse goblet cell hyperplasia |
| **$CD4^+$ T lymphocyte infiltration** | 0 | None |
| | 1 | Minimal |
| | 2 | Mild |
| | 3 | Moderate |
| | 4 | Abundant |

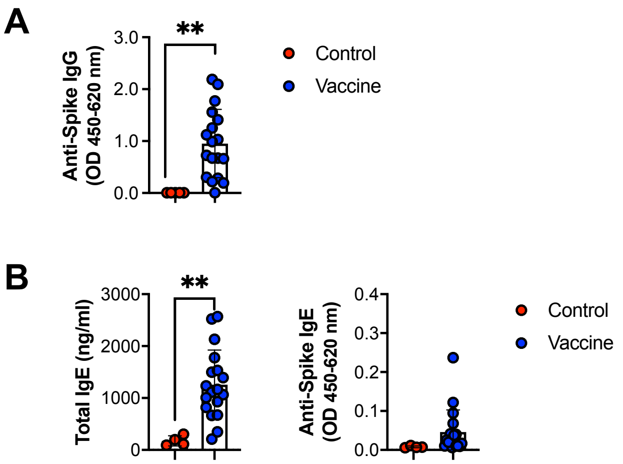


**Supplemental Figure S1: Spike-specific IgG and IgE production following Spike protein vaccination.**

(**A**) Spike-specific IgG production. (**B**) Total IgE and Spike-specific IgE production. Control n = 4, Vaccine n = 18. ** $p < 0.01$ by two-tailed student's *t* test. Data presented as Mean ± S.E.M..

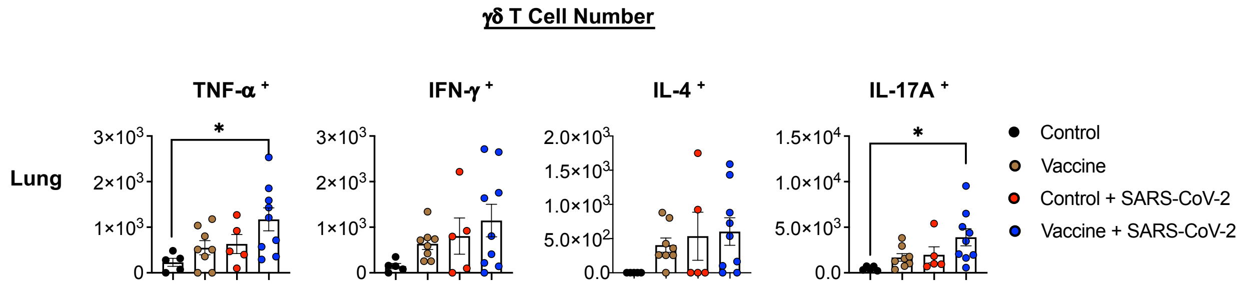


**Supplemental Figure S2: Effector cytokine production by lung γδ T cells.**

Samples in Figure 5 were also analyzed for γδ T cells. Numbers TNF-α, IFN-γ, IL-4 and IL-17A expression by viable γδ T cells in lung, lymph node and spleen are shown. Control n = 5; Vaccine n = 8; Control + SARS-CoV-2 n = 5; Vaccine + SARS-CoV-2 n = 9. *$p < 0.05$ by one-way ANOVA with multiple comparisons. Data presented as Mean ± S.E.M..
